# Understanding conformational dynamics from macromolecular crystal diffuse scattering

**DOI:** 10.1101/2021.02.11.429988

**Authors:** Parichita Mazumder, Kartik Ayyer

**Affiliations:** Max Planck Institute for the Structure and Dynamics of Matter, 22761 Hamburg, Germany; Center for Free Electron Laser Science, 22761 Hamburg, Germany; The Hamburg Center for Ultrafast Imaging, 22761 Hamburg, Germany

## Abstract

All macromolecular crystals contain some extent of disorder. The diffraction from such crystals contains diffuse scattering in addition to Bragg peaks and this scattering contains information about correlated displacements in the constituent molecules. While much work has been performed recently in decoding the dynamics of the crystalline ordering, the goal of understanding the internal dynamics of the molecules within a unit cell has been out-of-reach. In this article, we propose a general framework to extract the internal conformational modes of a macromolecule from diffuse scattering data. We combine insights on the distribution of diffuse scattering from short- and long-range disorder with a Bayesian global optimization algorithm to obtain the best fitting internal motion modes to the data. To illustrate the efficacy of the method, we apply it to a publicly available dataset from triclinic lysozyme. Our mostly parameter-free approach can enable the recovery of a much richer, dynamic structure from macromolecular crystallography.

---

X-ray crystallography remains the preeminent technique to determine the structures of macromolecules at atomic resolution. During a crystallography experiment, in addition to the bright and sharp Bragg peaks used for crystallographic analysis, one also observes weaker scattering at all angles, collectively termed diffuse scattering. The Bragg peak intensities encode information about the average electron density in the unit cell of the crystal, which in turn is used to generate a model of the average atomic positions and their variance (B-factors or ADPs) [1]. Part of the diffuse scattering is composed of trivial contributions from the Compton scattering, bulk solvent scattering, air scatter and other parasitic scattering from beam line components. But crucially, when the crystal is not perfectly ordered, it also contains photons scattered from the proteins themselves. For brevity, in the rest of the article, we will refer to this disorder-induced scatter as diffuse scattering, while keeping in mind that the other contributions must also be accounted for while preprocessing the data [2, 3].

If one assumes the disorder is not too large, the displacements of individual atoms are expected to follow a normal distribution (an assumption borne out by the success of the B-factor formalism in crystallographic refinement). Within this limit, the measured diffuse scattering is a function of the two-point correlation function of all atomic displacements. This means that one has direct access to equilibrium dynamics of proteins via correlations in their atomic displacements. In recent years, there has been an upsurge in interest in such data, driven by the desire to understand proteins as more than static objects since their equilibrium dynamics play a crucial role in their function. With modern detectors and X-ray sources, we can now measure these weak intensities with sufficient quality to enable interpretation [4].

Over the past three decades many models of motion have been proposed to describe the diffuse scattering including liquid-like motion [5–8], rigid-body motion [9– 12], lattice dynamics [3, 13] and normal mode analysis [14–17]. Another approach has been to examine the diffuse scattering predicted by molecular dynamics (MD) simulations of ever-larger systems [3, 18–21]. In the latter case, if MD predictions agree with the observed diffuse scattering, one can analyze the trajectories to gain insights into eqilibrium protein dynamics.

It has also been observed that agreement between experimental and calculated diffuse maps improve when motion due to coupled displacements of neighboring molecules in the lattice is included [22]. Recently, Meisburger, Case and Ando (MCA) reported a carefully processed dataset on triclinic lysozyme and analyzed it with a lattice dynamics normal-mode model [3], leading to the highest correlation coefficients with the experimental data yet. Unfortunately, these strong results still do not provide insights into functionally relevant dynamics of the proteins, since intermolecular interactions in a crystal are not likely to be relevant in the native environment. The long term goal of extracting a dynamical structure from macromolecular crystallography data requires a different approach, which we describe below.

In order to understand the conformational landscape sampled by crystallized proteins from their diffuse scattering, we employ three concepts which have not been utilized previously in the field. The first is to capitalize on the separation of diffuse scattering from long- and short-range correlations in reciprocal space. Secondly, we develop a Bayesian optimization pipeline to refine the dynamical parameters of our model which minimizes the number of computationally expensive calculations of the scattering from given parameters. Finally, we use restrained molecular dynamics to provide a basis for fitting against the measured diffuse scattering which better identifies the dominant modes in crystallized proteins.

We will expand upon each of these pillars as well as describing the dataset and system on which we will apply our approach in the results section. We show a significant improvement in the fit to the measured data upon inclusion of an optimized internal dynamics model. This analysis can also be extended to other systems and experimental datasets.

## RESULTS

### Short- and long-range correlations

The first property of diffuse scattering we exploit is that long-range correlations in displacements produce sharp intensity features and vice versa. As an extreme example, if the displacements are completely uncorrelated between atoms (correlation length is zero), the diffuse scattering is unstructured and contains no information apart from the average amount of disorder [23]. The other extreme is if the displacements of all atoms are fully correlated, in which case there is no disorder, and the “diffuse” scattering consists of just the Bragg peaks.

A special crossover happens when the correlation length increases past the unit cell size. Above this limit, if we assume that the crystal is homogeneous, then even though the atoms in each unit cell are displaced randomly, the probability distribution of the displacements described by the covariance matrix is the same for all unit cells. This means that the covariance matrix of atomic displacements has the symmetry of the crystal, and so too should the diffuse scattering produced by such long-range disorder, resulting in scattering in and around the reciprocal lattice points. This form of diffuse scattering has been observed and attributed to long range liquid-like correlations [22] and thermal acoustic phonon dynamics [3, 13]. In fact, such ‘halos’ around Bragg peaks are expected whenever intermolecular correlations monotonically decay with distance smoothly, as we expect if no special phonon modes are excited (see Methods section ‘Diffuse scattering from decaying intermolecular correlations’ for a mathematical justification).

If the dynamics of each unit cell are independent, then the diffuse scattering does not ‘see’ the reciprocal lattice at all. One example of this is uncorrelated rigid body translational (RBT) disorder [10] where the diffuse scattering is just the molecular transform weighted by the complement of the Debye-Waller factor. But this could also be coupled with conformational variations as described by simple models like liquid-like motion (LLM) which extend the RBT model to allow relaxation of strict rigidity when pairs of atoms are far apart. Such intramolecular correlations are the ones relevant to the structural flexibility of the molecule itself.

In general, when both intra- and inter-molecular correlations are present, their relative contributions to the diffuse scattering at a given **q** are strongly dependent upon its distance to the nearest reciprocal lattice point. In Fig. 1(a-c) we illustrate the reciprocal space separation of diffuse scattering from long- and short-range correlations with a toy model. Figure 1(a) shows the diffuse scattering from a crystal where the unit cell can be in one of two states (shown in inset) with random but equal occupancy and with some random translational disorder. This translational disorder is correlated across unit cells and is shown in Fig. 1(b) where only lattice points are shown and colored by their displacement magnitude. If one replaces each lattice point randomly by one of the two states, the crystal diffraction is shown in Fig. 1(c), where one can see the ‘halos’ from the distorted lattice on top of the smooth scattering from the two-state model. This kind of separation is derived from some basic assumptions in the Methods section ‘Diffuse scattering from separable correlations’.

**FIG. 1.**
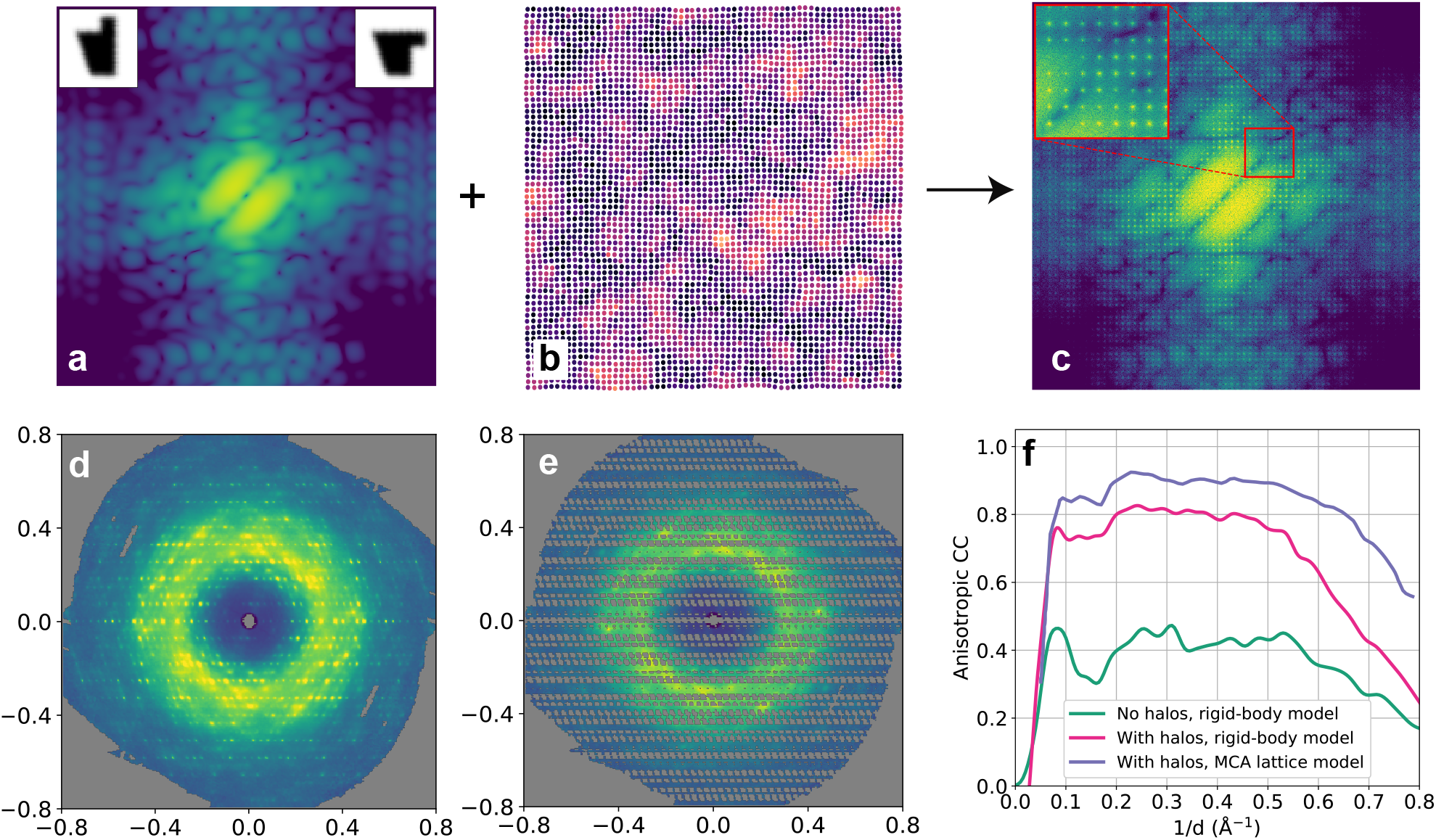
Toy model and sample system. (a) Diffuse scattering expected from an equally populated two-state model combined with rigid body translational disorder. The two states are shown in the insets. (b) Schematic of lattice dynamics of disorder (exponentially decaying correlations). Lighter colors represent lattice points further from their ideal positions. (c) Diffuse scattering when the unit cells are displaced according to the lattice in (b) and each unit cell is randomly in one of the two states. One can clearly see that other than the ‘halos’ around Bragg peaks (shown in the red box), the smooth diffuse intensity is the same as the model shown in (a). Both (a) and (c) intensities are shown on a logarithmic scale for clarity. (d) Slice through the experimental 3D diffuse intensity distribution, showing both sharp and smooth features. (e) Same slice as (d), but with the sharp ‘halos’ surrounding each Bragg peak masked out. (e) Pearson cross-correlation between a simple disorder model with rigid molecules and exponentially decaying intermolecular correlations and the experimental data with and without the halo features. Higher CC values obtained when the halo features are included contain the relatively trivial information that the ‘halos’ are bright and that they surround reciprocal lattice points. We also show the CC values obtained by MCA using their more sophisticated lattice model for intermolecular displacements.

Thus, in many cases, one can separate the scattering from inter- and intra-unit cell correlations in reciprocal space. The smooth diffuse scattering far from the Bragg peak ‘halos’ are sensitive to intra-unit cell or internal displacement correlations. This has been implicitly recognized earlier in attempts to “filter-out” these ‘halos’ to access the internal dynamics [2, 7]. We adapt an approach of masking out the halo features to avoid artifacts due to filtering and since our optimization algorithm can tolerate some missing data.

Another important point which one notes is that the long-range intermolecular correlations do have an effect on the scattering far from Bragg peaks. This is in the form of a rigid body translation component to the disorder within a unit cell. This is not to say that the molecules are rigid, rather that a significant part of the total disorder may be incorporated in a concerted motion due to lattice effects, but each molecule may be conformationally different on top of that.

### Model system

We apply our approach to the dataset collected by MCA on triclinic lysozyme crystals (PDB: 6o2h, CXIDB: 128) [3]. A central slice through their measured intensity distribution is shown in Fig. 1(d). In addition to being carefully analyzed in terms of background characterization and merging, the triclinic *P* 1 lattice also has the advantage of having a single asymmetric unit per unit cell, avoiding possible complications due to molecules being in different orientations. The authors observe that the dominant features were ‘halos’ surrounding Bragg peaks showing a 1*/* | *q−q*_Bragg_|^2^ intensity falloff, resembling the well-known behavior of thermal acoustic phonons. These are modes where the displacement of one molecule induces displacements in its neighbors, resulting in longitudinal waves with long correlation lengths compared to the lattice constants. MCA analyze these ‘halos’ to reconstruct the connectivity network of the crystal while assuming the molecules themselves to be rigid. A small improvement is observed by allowing the molecular structure to vary according to a previously predicted normal mode.

Since our primary interest lies in understanding the internal dynamics, we mask the regions of reciprocal space close to reciprocal lattice points and use the rest to optimize our dynamics model for a single protein. This has the additional computational benefit of needing to only simulate the MD trajectory of a single molecule, rather than a super-cell [3, 21], making it more computationally accessible as a technique. Figure 1(e) shows a slice through the masked intensities for the lysozyme model dataset. One effect of this masking is that the correlation coefficient between the measured and predicted data is lower than if the ‘halos’ were retained. This is due to two reasons, first that the measured intensities in this region are weaker, and hence noisier. The other, more important reason is that any model with ‘halos’ will have a relatively high Pearson correlation coefficient (CC) simply because it predicts high values near the Bragg peaks.

This is highlighted in Fig. 1(f) where the CC is calculated with and without masking. A rigid-body translational disorder model is used for the intra-cell dynamics, whose predicted scattering is just the *q*-weighted molecular transform. The anisotropic CC is around 0.4 for most of the resolution range when comparing against masked data. However, the introduction of even a simple common multiplicative halo function increases the CC to 0.75 (see Methods section for estimation of ‘halos’). The “lattice” model with more sophisticated ‘halos’ from MCA is shown for comparison.

### Refinement framework

We now turn our attention to determining what internal variability modes best describe the diffuse scattering far from the Bragg ‘halos’. In the rest of this study, we will focus on this masked diffuse scattering since this limits us to correlations at length scales much less than the lattice constants.

As discussed above, within the harmonic approximation, this diffuse scattering is a function of the pair correlation tensor 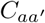 between atoms *a* and *a*′ within the molecule. Unfortunately, this mapping is not invertible since the number of independent measurements, which is of the order 𝒪 [(*d*_protein_*/d*_resolution_)^3^]is much smaller than the number of unknowns, 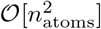 for protein crystals. This means that there should be many 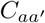 solutions which generate the same diffuse scattering. But almost all those solutions will be highly unphysical, corresponding to motions which have a very high free energy cost. Thus, one would like to optimize the 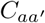 matrix within a subspace of physically reasonable distortions of the molecule.

We use molecular dynamics (MD) to generate this subspace. Any MD trajectory can be converted to a (3*N×*3*N*) 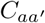 matrix by calculating the covariance of the displacements of all 3 components of all atoms over the trajectory. This matrix can then be diagonalized to calculate the so-called “essential dynamics” of the trajectory [24]. It has been observed that most of the 3*N* modes generated as the eigenvectors of this diagonalization are occupied with low amplitude, collectively giving rise to effects similar to uncorrelated motion of the atoms. There are only a few essential modes which describe the low free-energy parts of the motion in the trajectory.

Now the trajectory itself can be used to predict the diffuse scattering, but this has had limited success predicting scattering features [20]. We adopt the hybrid approach of relying on MD to generate the basis of likely modes, but refine the weights of the modes against the measured diffuse scattering.

The first step in performing this refinement is the forward calculation of the diffuse scattering given a set of modes and their relative weights. By *weight*, we mean the amplitude of random oscillations along that mode. If the average position and structure of the molecule is represented by the 3*N*-dimensional vector ⟨**r** ⟩ and the *i*-th displacement mode is Δ_*i*_, the distorted molecule has the coordinates

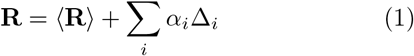

where *α*_*i*_ is a scalar which is the magnitude of the distortion along mode *i*. The weights to be optimized are the standard deviations of the random numbers *α*_*i*_. In order to calculate the diffuse scattering, a large number of randomly distorted molecules are generated to make an ensemble of structures corresponding to a given set of mode weights. The Guinier equation is then used to calculate the diffuse scattering as follows

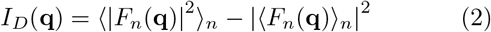

where *F*_*n*_(**q**) is the molecular transform of the *n*-th distorted molecule. One immediate feature of this kind of calculation is that it is noisy, on top of the noise in the measured data. This Monte Carlo sampling of the molecular distortions means that in order to reduce the noise in the calculation, one requires a large number of samples, making it expensive to evaluate.

An optimization method purpose-built for such problems is Bayesian global optimization (BGO) [25–28]. In this method, a noise-aware surrogate function is fit against the measurements. This surrogate function is fast to evaluate and so can be optimized using standard methods. The method then also predicts the optimal choice for the next sample. Thus, it is ideal for situations where one has a black-box, expensive to evaluate, noisy forward function without gradients. The goal is not just to optimize the objective function, but also to minimize the number of evaluations of our expensive function. The objective function we employ is the anisotropic cross-correlation coefficient in a relevant resolution range (Eq. 13).

A flow chart of the whole reconstruction algorithm is shown in Fig. 2(a). Figure 2(b) shows the noise in the objective function for a particular set of weights as a function of the number of Monte Carlo samples. To illustrate the method, we show the results of the algorithm for a 2-parameter optimization of the liquid-like motion parameters *σ* (rms of displacement) and *γ* (correlation length). Figure 2(c) shows the convergence of the best objective function with iteration number. Figure 2(d) and (e) show the reconstructed surrogate function as a function of *σ* and *γ* and the sampled points respectively. Details of the optimization pipeline are given in the Methods section.

**FIG. 2.**
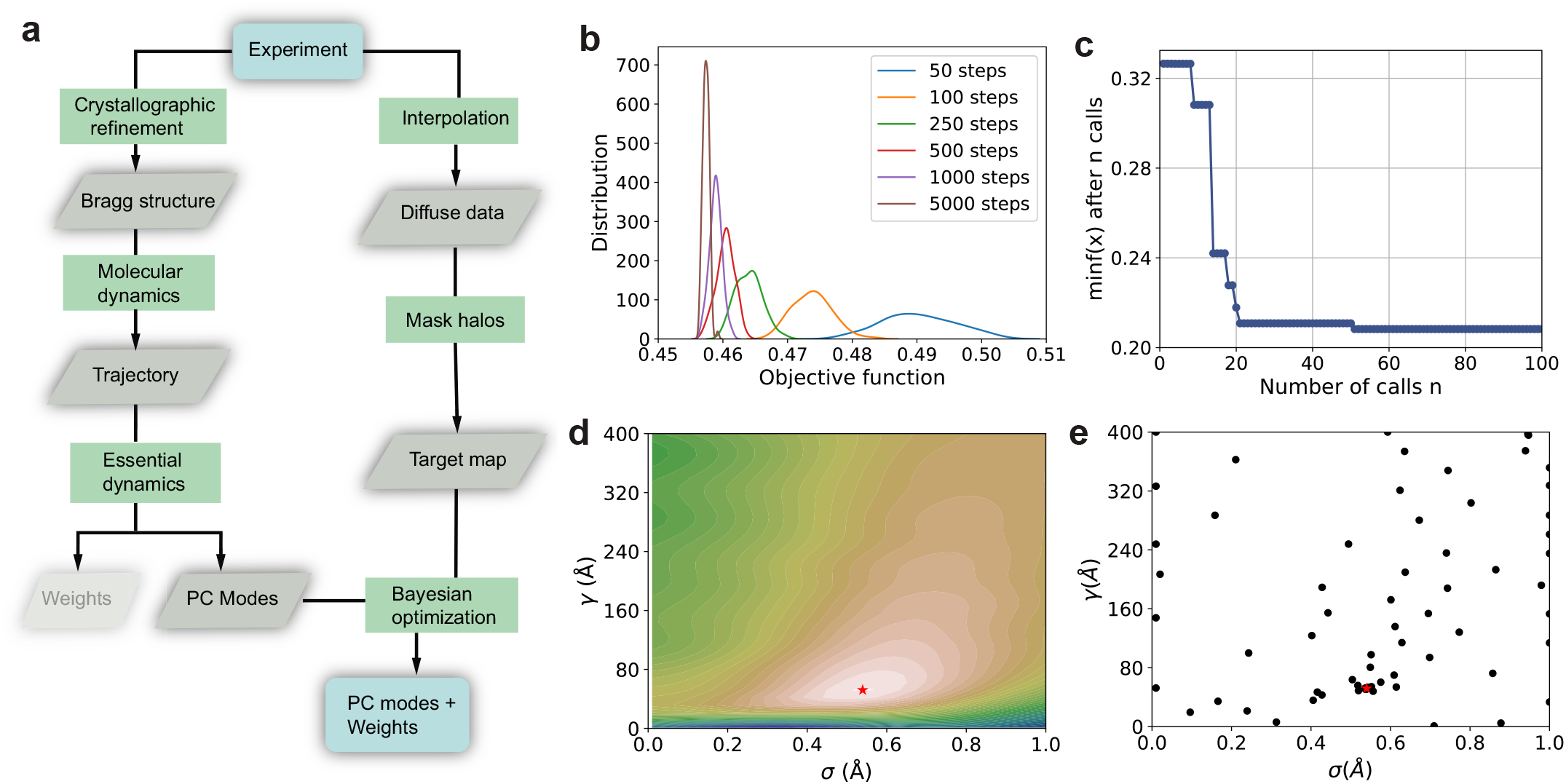
Reconstruction pipeline. (a) Flow-chart of pipeline to estimate principal component (PC) internal dynamics modes and their weights. Molecular dynamics is used to estimate energetically favorable internal modes and their weights are determined using Bayesian optimization by fitting against the measured diffuse scattering data. (b) Distribution of objective function values from different Monte Carlo runs with the same parameters. The width of this distribution is used to parameterize the Gaussian process regressor used in the Bayesian optimization. (c) Convergence plot showing the evolution of the best objective function with iteration number. This is the output for a 2-parameter optimization of liquid-like motion parameters, *σ* (rms displacement) and *γ* (correlation length) for demonstration. (d) Reconstructed surrogate objective function. (e) Points sampled in *σ − γ* space, with denser sampling close to the optimum.

### Position-restrained molecular dynamics

In order to apply the framework designed above, we need a basis set of dynamical modes which can describe the covariance matrix of atomic displacements. We generate this basis from a molecular dynamics simulation using the ideas of ‘essential dynamics’ [24]. Briefly, one generates a molecular dynamics (MD) trajectory, from which the displacement covariance matrix is calculated. This matrix is diagonalized to generate eigen-modes and one observes that, usually, only a few modes account for the bulk of the correlated motion.

Proteins in crystalline environments have more constraints to their motion than those in solution due to the presence of the neighboring molecules. One way to account for this is to simulate a super-lattice with multiple unit cells of the crystal [3, 21], which can be computationally expensive. We choose an alternative strategy where we simulate a single molecule but apply additional positional restraints to all non-hydrogen atoms. The strength of the restraining force is a function of the crystallographically refined B-factor of that atom and the effective B-factor of the atom from an unrestrained MD simulation. If the root mean squared fluctuation (RMSF) of an atom from an MD trajectory after rigid body alignment is *σ*, the effective B-factor is 4*π*^2^*σ*^2^.

For an isolated atom simulated at temperature *T*, a force constant of *F* results in a B-factor of *k*_*B*_*T/F* [29]. One can use this to estimate the effective force constant on each atom *i* from the intrinsic MD force fields,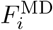. The total restraining force to produce the measured B-factors should be 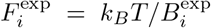. For our model system, unrestrained MD almost always produces much higher atomic fluctuations than the experimental data. Thus, one can calculate an additional restraining force, 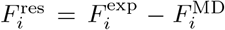. But this is true only in the case where atomic displacements are independent. The presence of structured diffuse scattering means that displacements are correlated and so restraining just one atom reduces the fluctuations of many others by suppressing otherwise favored modes of motion.

In our system, we found a reasonable agreement between restrained-MD and experimental B-factors by using force constants 200 times lower than those predicted by the above calculation (see Fig. S1 for details) The agreement was not perfect, but is sufficient to significantly improve the correlation with the experimental data.

### Application to triclinic lysozyme

The experimental data was masked as described above in the section on short- and long-range correlations by masking out a spherical region around each reciprocal lattice point whose radius was 3*/*7-th of the inter-peak spacing. This region was spherical in *hkl* space, but ellipsoidal in isotropic reciprocal space since the lattice was triclinic. A slice through the masked intensities is shown in Fig. 1(b).

In order to generate an ensemble of structures from which to calculate the principal component modes, we performed a 1 µs simulation of a single solvated lysozyme molecule with average structure given by the crystallographic structure (PDB: 6o2h). Position restraints were applied as described above in the restrained MD section and the correlation coefficient (CC) with the masked diffuse scattering of the trajectory itself is shown in Fig. 3(a). The covariance matrix of fluctuations of the positions was diagonalized to generate the principal component dynamical modes and their relative weights.

**FIG. 3.**
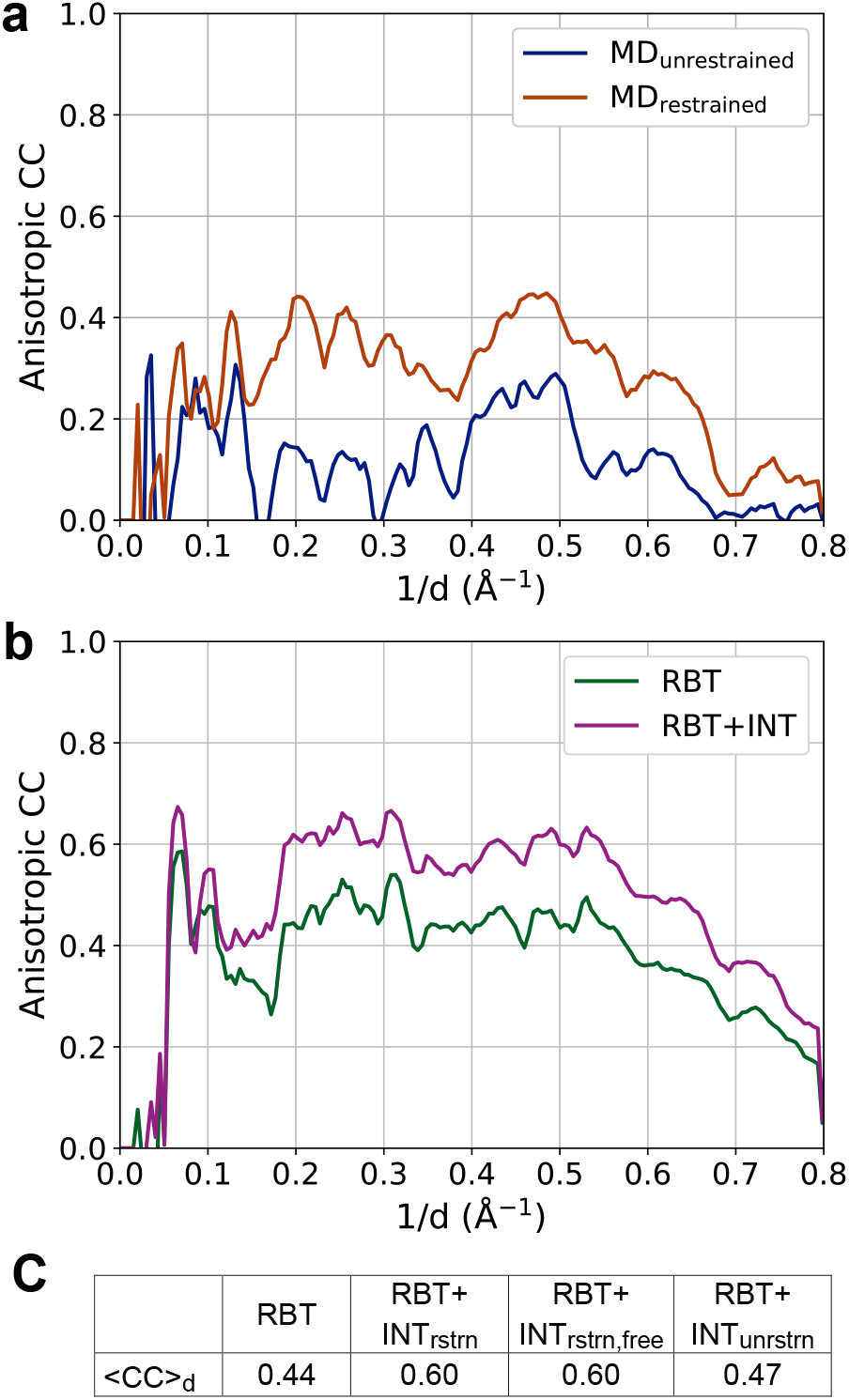
Cross-Correlation between masked experimental data and simulated diffuse intensity. (a) Diffuse scattering calculated from unrestrained and restrained MD trajectories. (b) Different short-range models: rigid-body translation (RBT) and RBT + internal modes (RBT+INT) with RBT *σ* = 0.46Å. (c) Summary of average cross-correlation (CC) in the resolution range d, 3.3Å−1.6 Å for different models.

This data was compared against the masked experimental data. Various Bayesian optimization runs were performed, and the results are summarized in Fig. 3(b). The base model was isotropic rigid body translation (RBT), which generates an average CC of 0.44 in the (3.3 Å-1.6Å) resolution range. The weights of the principal component modes were optimized along with rigid body translation as described above. That resulted in a significant increase in the average CC in the same resolution range to 0.60 with the inclusion of this internal dynamics (RBT+INT). With the inclusion of liquid-like motion (LLM) and no internal dynamics, the average CC in that range was 0.60 (RBT+LLM). The optimization of internal dynamics in the presence of liquid-like motion increased the CC to 0.62 (RBT+LLM+INT) (data not shown for clarity). The more modest improvement is partially explained by the effect that LLM-like effects can result from many equally weighted modes, each of whom involve the concerted motion of a part of the molecule.

For validation of our INT model we performed Bayesian optimization with top 10 unrestrained MD modes which gave a CC of 0.44 in the same resolution range, which is the same as what we get with RBT only, showing that the wrong choice of basis does not improve the fit to the data even though additional parameters are refined. We also performed an optimization by excluding the Brillouin zones around 10% of the Bragg peaks and calculated the CC_work_ and CC_free_ in the standard way (see Fig. S2). The average CC_work_ was 0.61 while CC_free_ calculated just from the excluded voxels in the dataset was 0.60.

Figure 4 illustrates the most prominent mode found in the (RBT+INT) optimization. This mode describes motions between conformations which are present in crystallized lysozyme Fig. 4(a). To describe the internal motion of this mode, the displacement of the C_*α*_ backbone is shown in Fig. 4(b). The *β*-lobe (THR40 to CYS94) shows higher motion compare to the *α*-lobe (ARG5 to SER36 and ILE98 to LEU129). The displacement is low for active site GLU35 and ASP52. Some internal deformation is observed in the *β*-lobe Fig. 4(c-d). Around the hinge region, ASN39 shows significant movement. This and other prominent modes are illustrated in Supplementary Movie S1.

**FIG. 4.**
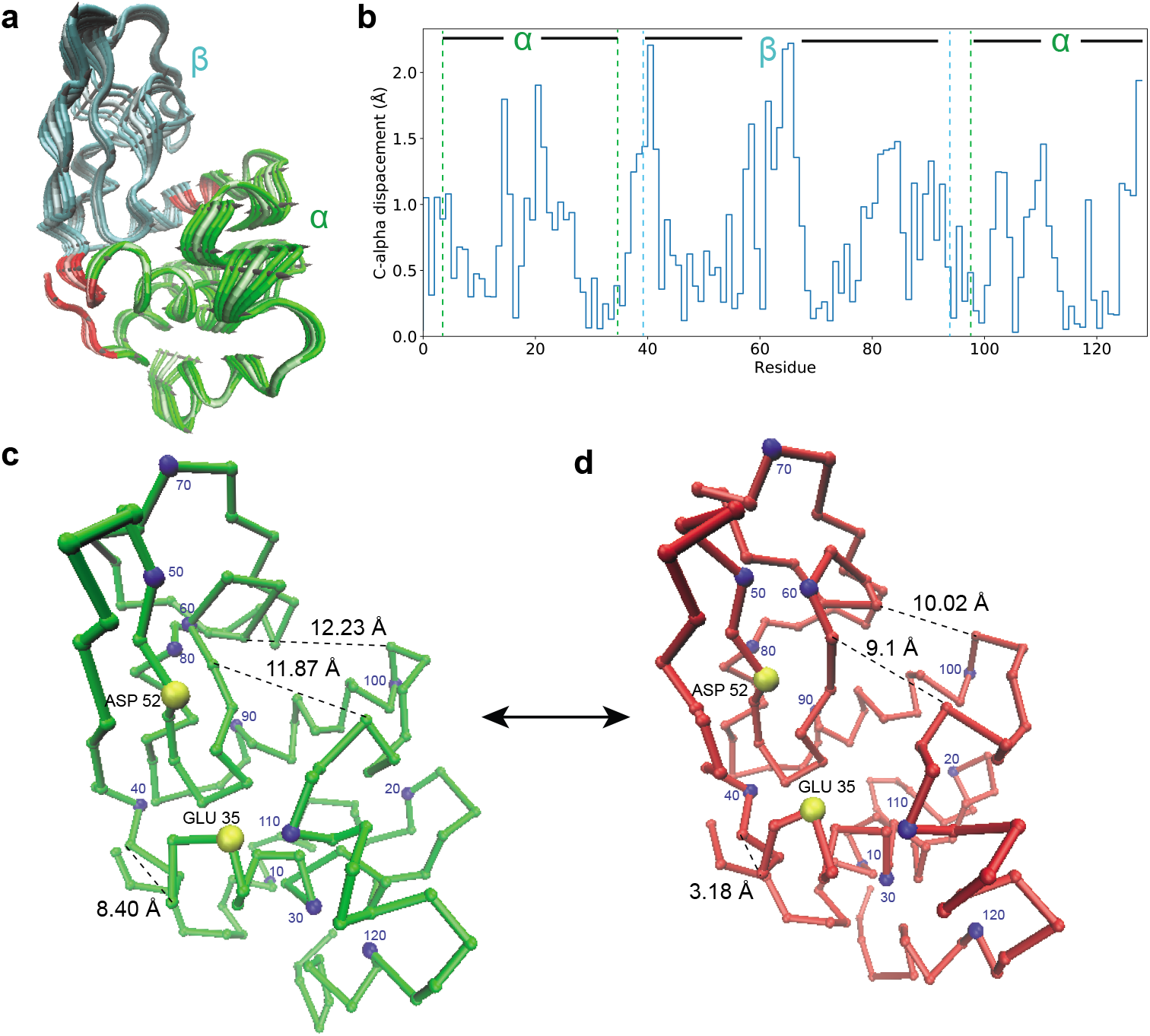
Visualization of dominant mode of motion. (a) Conformations along the dominant principal mode. The mode is shown in both directions black arrows. (b) C_*α*_-atom displacement for this mode. The *α* and *β* lobe atoms are shown within cyan and green dotted lines. (c-d) C_*α*_ atoms displacement in two different conformations shown in green(far) and red(close).The distance between ASN37-ASN39, TRP63-ASP101, ASN59-ALA107 are shown to understand the motion. Every 10th C_*α*_ atom is colored in blue. The active site residue GLU35 and ASP52 shown in yellow. See Movie S1 for a visualization of this and other modes with non-negligible weights.

## DISCUSSION

Proteins are inherently dynamic objects, with their conformational variation often playing a crucial role in their function. While conventional crystallography can routinely provide high resolution static structures, they provide no direct information about correlated motion within the molecule [4]. Diffuse scattering, which is always present but usually discarded, is affected by the pair correlation of the displacements, and hence is strongly influenced by whether different atoms are displaced synchronously or asynchronously and to what extent.

In this article, we demonstrate a method to fit the internal dynamics of a protein to its diffuse scattering. The first step in this process is the partitioning of the effect of conformational motions from lattice dynamics in reciprocal space for homogeneous crystals. This separation can be detected in the form of e.g. ‘halos’ around reciprocal lattice points, and the signal far from such lattice-induced features is dominated by internal structural variation of the protein within a unit cell. Once the relevant part of the experimental data is identified, we use molecular dynamics to obtain a basis space for dynamical modes whose weights are optimized using a Bayesian optimization procedure.

As a proof of principle, we apply this method to the diffuse scattering from triclinic lysozyme crystals [3]. Our first observation is that the basis space generated from the MD simulation of isolated proteins in solution does not fit the data well, suggesting that the dominant modes in solution are poorly represented in the crystalline environment. Better agreement with the data was found by applying additional restraining force fields which more closely align the fluctuations in the atomic positions to the experimental B-factors. The optimized dominant modes improve the correlation coefficient with the experimental data to 0.60 from the 0.45 value obtained from a rigid body model of the protein. Note that we have to include rigid body motion along with the internal modes to obtain good agreement, which may explain the poor agreement observed in previous studies incorporating only internal dynamics which hold the center-of-mass of the molecule fixed [22]. The strongest conformational mode in the crystalline environment predicted from the optimization is shown in Fig. 4.

Apart from the initial analysis of the ‘halos’, the whole procedure is mostly parameter-free. With further refinements to the pipeline, we envision the analysis of such internal dynamics to be routinely performed on crystallographic datasets to create a dynamic model of the molecule, and not just the coordinates plus B-factor static picture we currently extract. One future direction of improvement is a better scheme for restrained MD simulations such that the first few modes are the most relevant to the crystalline dynamics. At the other end, the optimization scheme could also be improved to allow us to increase the number of modes which can be simultaneously optimized. Of course, this also requires improvements in the collection and processing of the experimental data, especially with regards to reducing and characterizing the background scattering.

## METHODS

### Diffuse scattering from decaying intermolecular correlations

Following the convention of [23], the position of atom *a* in cell *c* is **r**_*ac*_ = **r**_*c*_ + **r**_*a*_ and the total intensity from a disordered crystal is given by

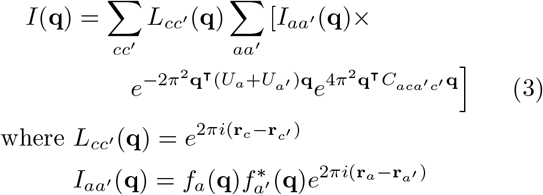

where *U*_*a*_ is the 3×3 variance matrix of atomic displacements, also called the anisotropic displacement parameter (ADP) and 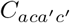 is the covariance matrix of the displacements of atoms *ac* and *a*′*c*′.

Here we analyze the diffuse scattering when the intermolecular correlations are non-negative and smoothly decaying as a function of the separation between the unit cells. For simplicity the molecules themselves are assumed to translate as rigid units. The next subsection discusses the case when this latter assumption is not true. This generalizes the derivation of scattering for liquid-like motion in [23]. From the above mentioned assumptions, let 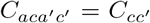 be a function of only 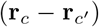 for all *a, a*′. Then, Eq. 3 becomes

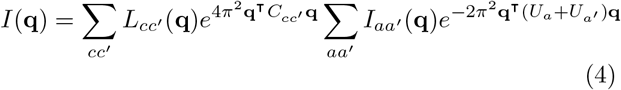

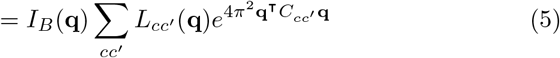

where *I*_*B*_(**q**) is just the Bragg intensity distribution obtained from the Fourier transform of the average unit cell. The exponential in the weighted lattice sum can be expanded to give

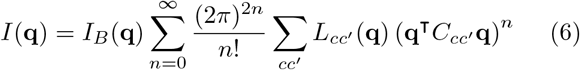

Expanding the 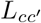 term and including the assumption that 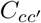 is only a function of 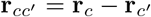, we can see that the sum only depends upon the separation 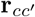. This allows us to insert the integral term 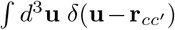 and get

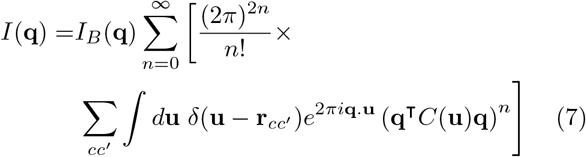

The integral is now just a Fourier transform integral,

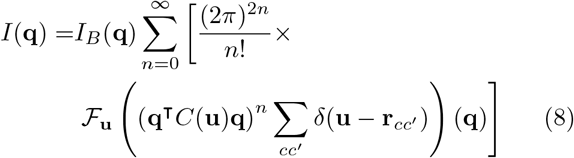

Where ℱ represents the Fourier transform operation. From the convolution theorem,

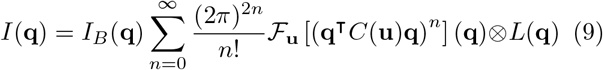

where *L*(**q**) is the lattice sum generated by the Fourier transform of an ideal lattice. The *n* = 0 term corresponds to the Bragg intensities while the other terms contribute to the diffuse scattering. This series represents a sum of broadened reciprocal lattices where the higher order terms are weighted towards high resolution due to the *q*^2*n*^ dependence. *C*^*n*^(**u**) gets sharper as *n* increases which has the reverse effect on its Fourier transform, broadening the convolution kernel for higher *n*.

Thus, we get the result that when lattice interactions result in correlations which decay with inter-molecular separation, the diffuse scattering is given by a series of broadened Bragg peaks (‘halos’) where the peaks get broader at higher resolution.

### Diffuse scattering from separable correlations

In this section, we analyze the diffuse scattering caused by displacement correlations that can be separated into intra- and inter-molecular terms. Thus, the covariance matrix 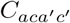 between atoms *ac* and *a*′*c*′ can be described as the sum of two terms

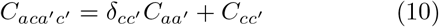

In this formulation, atoms within a molecule have correlated displacements governed by 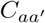 but the interac tion between molecules is governed purely by 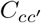 for all atoms in the molecule. Physically, this corresponds to different energy scales or mechanisms for internal interactions versus those for lattice interactions.

Then Eq. 3 becomes

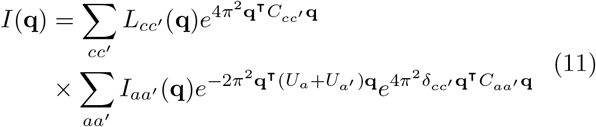

Separating the two cases when *c* = *c*′ and *c ≠ c*′ and then completing the sum, we get

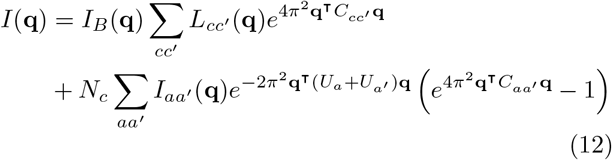

The first term results in the ‘halos’ as discussed in the previous subsection while the second term is independent of the lattice periodicity and only depends on the internal dynamics of the molecule, 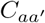.

The target smooth diffuse scattering map which corresponds to the short-range correlations, was generated from experimental data by masking out a spherical region around each reciprocal lattice point(‘halos’).

### Simple model for estimation of ‘halos’

To illustrate the effects of including ‘halos’ on the CC metric, a simple uniform ‘halo’ model was estimated directly from the experimental data. The ‘halos’ were simulated by broadening the Bragg intensities by a ‘halo’ kernel, whose shape was estimated as described below.

The signal in each Brillouin zone was averaged, leading to a 7 *×*7 *×*7 voxel volume. The minimum value was subtracted and the peak was exponentiated by a variable exponent *α* and tiled out to generate a lattice-sum function. This function was multiplied to the rigid-body diffuse scattering to generate the predicted intensities with ‘halos’. A one-dimensional scan was performed to optimize the *α* parameter, leading to a value of 3.8 used in estimating the CC values shown in Fig. 1(c).

### Molecular dynamics simulations

MD simulations were performed using GROMACS 2019.4 [30] using OPLS-AA/L all-atom [31] parameters for protein and OPLS-2009L parameters for nitrate ions [32]. The starting structure was generated using coordinates from PDB ID 6o2h (conformation A for lysozyme with seven nitrate ions, one chloride ion and crystal waters). Next, this model was solvated with a TIP3P water box of dimension 60*×*67*×* 74 Å and neutralized with one chloride ion. After minimization (steepest decent), the model was equilibrated for 1 ns with position restraints on protein atoms. Then, two sets of simulation runs were performed in NPT conditions to keep the temperature and pressure stable around 295 K and 1 bar with 2 fs time steps. For unrestrained-MD, we gradually reduced the position restraints on protein atoms to zero over 0.2 µs and then recorded coordinates every 10 ps for 1.7 µs.

For restrained-MD, we applied positional restraints on protein atoms to the equilibrated model (see Fig. S1 for details) during the 1 µs production run to maintain the experimental B-factors.

### Diffuse scattering calculation

For diffuse scattering calculation from internal dynamics, the trajectory was fitted with respect to the backbone of the starting configuration, comprising the last 150000 frames for unrestrained-MD and last 90000 frames for restrained-MD. To generate the essential modes of motion from MD trajectory we calculated a (3*N ×* 3*N*) covariance matrix of the displacements of all 3 components of all N atoms over the trajectory. This matrix was then diagonalized to calculate the essential modes of the trajectory. We used VMD, NMWiz to visualize the modes [33, 34].

For a given set of coordinates or mode variance vector (or covariance matrix), the diffuse scattering was calculated using the Guinier equation (Eq. 2, in results ‘Refinement framework’ section).

### Bayesian global optimization (BGO)

We implemented Bayesian optimization with Gaussian process regressors using the `scikitoptimize`Python package (v0.8)[35]. Since the goal was to maximize the correlation between experimental and simulated diffuse intensity, the objective function was defined as,

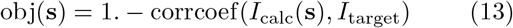

where the correlation was calculated after subtracting the radial averages of both terms and was averaged over the resolution range of 3.3-1.6 Å.

The larger the number of Monte Carlo simulation steps used to generate *I*_calc_, the lower and less noisy the value of the objective function (see Fig. 2(b)). To compromise between speed and precision, we used 1000 samples during the optimization while the final CC was calculated using 5000 samples. We used BGO to find the minimum of the objective function over the range of one or multiple input parameters, **s**, in as few iterations as possible. The Gaussian process surrogate model gives an estimate of the objective function which can be used to direct future sampling. For acquiring more samples we used `gp hedge`acquisition function to minimize over the Gaussian posterior distribution with the `lbfgs`acquisition optimizer. A noise level of 1*×*10^−7^ was used during optimization with `n_restarts_optimizer`= 2. We performed various optimization runs with respect to masked diffuse data with different short-range models, e.g., 1-parameter optimization of rigid-body translation (RBT) *σ*, 2-parameters optimization of the liquid-like motion (LLM) parameters *σ* and *γ*, 10-parameters optimization of internal modes weights along with RBT *σ* = 0.46 Å (see Fig. S3) etc. For internal modes optimization we ran BGO 128 times from random initial starting estimates. We noted that excluding some failed runs, most of the optimized **s** values could be clustered around a single point. For convenience, we estimated the best fit internal mode weights by calculating the median value of the results from the 128 runs.

## Data availability

The experimental data used in this study were previously published on the CXIDB (ID: 128). The molecular dynamics trajectories and optimization outputs are available from the corresponding author upon reasonable request.

## Code availability

The code used to perform the calculations here can be found at https://github.com/AyyerLab/diffuser.

## ACKNOWLEDGMENTS

This work is supported by the Cluster of Excellence ‘CUI: Advanced Imaging of Matter’ of the Deutsche Forschungsgemeinschaft (DFG) - EXC 2056 - project ID 390715994. The computations in this work were partially performed at the Max Planck Computing & Data Facility. We thank Henry N. Chapman and Thomas J. Lane for valuable comments on a draft of this article.

## Author contributions

K.A. conceived the project. K.A. and P.M. wrote the analysis program. P.M performed the molecular dynamics simulations and the analysed the output. K.A and P.M. wrote the manuscript.

