## Supplementary Information for "Understanding conformational dynamics from macromolecular crystal diffuse scattering"

### RESTRAINED MOLECULAR DYNAMICS B-FACTORS

An additional restraining force was calculated as $F_i^{\text{res}} = F_i^{\text{exp}} - F_i^{\text{MD}}$ for each atom $i$, where $F_i^{\text{MD}}$ is the effective force constant of the natural forces on that atom.

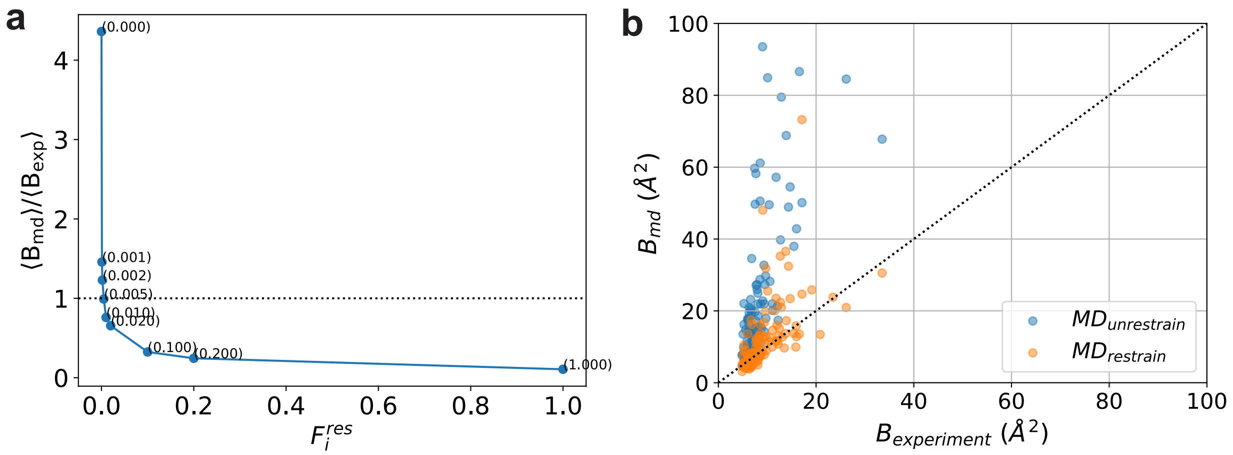


FIG. S1. Restrained MD. (a) Ratio of average B-factor of C-alpha atoms obtained from restrained MD and experiment. $\lambda$ values are shown in parentheses. (b) Scatter plot of isotropic crystallographic B-factors with those from restrained ($\lambda$= 0.005) and unrestrained MD runs. Even with atom-wise positional restraints, the correspondence is not perfect.

To get a reasonable agreement between restrained-MD and experimental B-factors we performed several sets of restrained-MD for 10 ns with an applied restraining force on each atoms, $F_i^{\text{app}} = \lambda F_i^{\text{res}}$. We found a reasonable agreement at $\lambda = 0.005$ (see Fig. S1(a)) and we continued this restrained-MD run for 1 µs to generate the trajectory for further analysis. Figure S1(b) shows a scatter plot of the atomic B-factors before and after applying additional restraints.

### VALIDATION OF REFINEMENT MODEL

We performed two validation tests for our internal motion model to eliminate over fitting. For first validation we performed Bayesian optimization(BGO) with top 10 unrestrained MD modes combine with RBT $\sigma$ 0.46 Å. The measured diffuse intensity was not a good fit with masked experimental data. We observed similar cross-Correlation as the measured diffuse

*

intensity from RBT only model Fig. S2(a). But for top 10 restrained MD modes we found significant improvement (Fig. 3(b)). Secondly we ran BGO by excluding the Brillouin zones of 10% of the Bragg peaks and calculated $CC_{work}$ and $CC_{free}$ Fig. S2(b). The similar $CC_{work}$ and $CC_{free}$ shows how our model is not over-fitting the data.

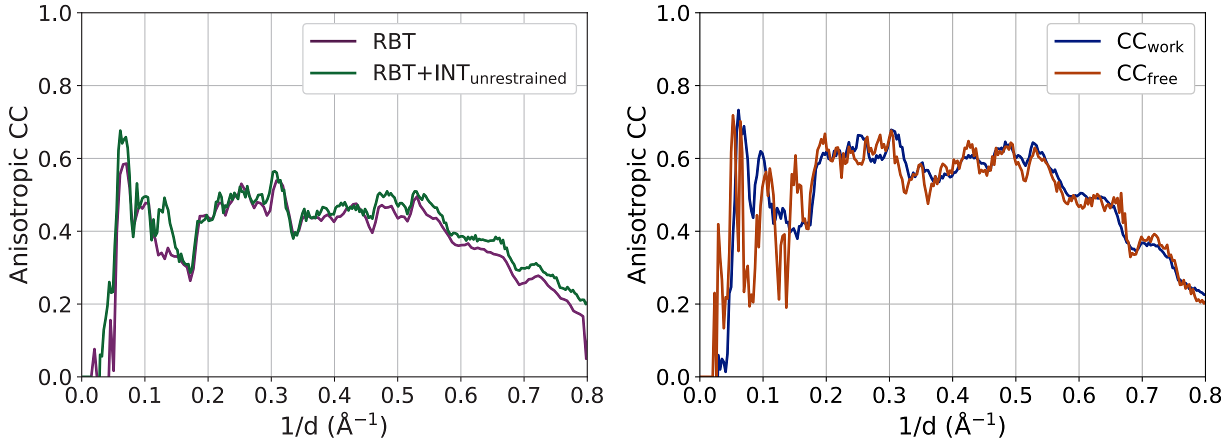


FIG. S2. Cross-Correlation between experimental data and simulated diffuse intensity. (a) Diffuse scattering calculated from rigid-body translation (RBT) and RBT + internal modes from unrestrained MD ($RBT+INT_{unrestrained}$) with RBT $\sigma = 0.46$ Åoptimized against masked experimental diffuse data. (b) $CC_{work}$ and $CC_{free}$.

### CONFIGURATION FILE

Figure S3 shows the configuration file used with the *diffuser* package for the final optimzation run. Operating instructions for the package are supplied with the source code at `https://github.com/AyyerLab/diffuser`.

### SUPPLEMENTARY MOVIE S1

Movie showing the conformational modes corresponding to highest weighted 6 of the top 10 modes obtained by diagonalizing the restrained MD trajectory. The numbers correspond to the weights of the modes scaled to the dominant mode 5. This mode is also described in Fig. 4.

```
[files]
topo_fname = lyso.tpr
traj_fname = protein900ns_fcby200_fit.xtc
pdb_fname = 6O2H.pdb
selection_string = protein and not altloc B and not altloc C
vecs_fname = md_fcby200_100vecs.h5

[parameters]
size = 315
res_edge = 1.26048
rot_axis = 2
sigma_deg = 0
sigma_vox = 0.46
num_steps = 1

[optimizer]
vecs_fname = md_fcby200_100vecs.h5
itarget_fname = intens_7_iso_cube_3.h5
output_fname = lyso_mdfcby200_rbfixed_6o2h_10vecs_aniso_hkl.pkl
num_steps = 1000
num_vecs = 10

diag_bounds = 0 2
calc_anisotropic_cc = yes
q_range = 0.3 0.6
```

FIG. S3. Configuration file used for the final optimization run to obtain the weights of the top 10 modes of the restrained MD trajectory. Full paths to various file names have been truncated for clarity.
